# Rabies-induced behavioural changes are key to rabies persistence in dog populations: investigation using a network-based model

**DOI:** 10.1101/554451

**Authors:** Victoria J. Brookes, Salome Dürr, Michael P. Ward

**Affiliations:** Sydney School of Veterinary Science, The University of Sydney, Camden, Australia; Veterinary Public Health Institute, University of Bern, Liebefeld, Switzerland

**Keywords:** Rabies, simulation modelling, canine, social network analysis, vaccination

## Abstract

Canine-rabies was endemic pre-urbanisation, yet little is known about how it persists in small populations of dogs typically seen in rural and remote regions. Our objectives were to simulate rabies outbreaks in such populations (50—90 dogs) using a network-based model, and investigate the influence of rabies-induced behavioural changes.

Such changes – increased bite frequency and either roaming or paralysis that increased the number or duration of contacts, respectively – were essential for disease propagation. Spread occurred in approximately 50% of model simulations and in these, a very low case rates (2.0—2.6 cases/month) over long durations (95% range 20—473 days) were predicted. Consequently, disease detection is a challenge, risking human infection and spread to other communities via dog movements. Even with 70% pre-emptive vaccination, spread occurred in >30% of model simulations (in these, median case rate was 1.5/month with 95% range of 15—275 days duration).

We conclude that the social disruption caused by rabies-induced behavioural change is the key to explaining how rabies persists in small populations of dogs. We predict that vaccination of substantially greater than the recommended 70% of dog populations is required to prevent rabies emergence in currently free rural areas.

## 1. BACKGROUND

Canine rabies is an ancient disease that has persisted in dog populations for millennia – well before urbanisation (1). Increased understanding of rabies spread in communities with relatively small populations of dogs – such as those in rural and remote areas –could give insights about rabies persistence in non-urban areas, as well as inform prevention and control strategies in such regions.

Rabies virus is neurotropic and clinical manifestations of canine-rabies can be broadly classified as the dumb form (characterised by progressive paralysis) and the furious form (characterised by agitated and aggressive behaviour; (2, 3). Although the mechanisms of rabies-induced behavioural signs are poorly understood (4) pathogen-influenced changes in host behaviour can optimise pathogen survival or transmission (5). We hypothesise that rabies-induced behavioural changes promote rabies transmission in dog populations by influencing social network structure to increase the probability of effective contact, and that this might be important to enable rabies to spread in rural and remote regions.

Since 2008, rabies has spread to previously free areas of Southeast Asia. Islands in the eastern archipelago of Indonesia, as well as Malaysia are now infected (6-9). Much of this regional spread of canine-rabies has occurred in rural and remote areas. Oceania is one of the few regions in the world in which all countries are rabies free. Recent risk assessments demonstrate that Western Province, Papua New Guinea (PNG) and northern Australia, are at relatively high risk of a rabies incursion (10, 11). Dogs in communities in these regions are owned and roam freely; population estimates in such communities are often low; for example, median 41 dogs (range 10—127) in Torres Strait communities (*pers comm* Queensland Health) and median 100 dogs (range 30—1000) in Western Province Treaty Villages (*pers comm* Australian Commonwealth Department of Agriculture and Water Resources). Canine rabies might have a low probability of maintenance in domestic dogs in these communities due to their small population sizes, but if continued transmission occurs – particularly over a long duration – then spread to other communities or regional centres and regional endemicity might occur.

GPS telemetry data from Torres Strait dog populations have recently been collected (12). Such data has been used to describe contact heterogeneity in animal populations, and has been used in models to provide insights about disease spread and potential control strategies (13-15). The effect of contact heterogeneity on disease spread is well-researched and models can provide useful insights about disease control strategies in heterogeneously mixing populations (16-18). Most recently in the context of rabies, Laager et al. (19) developed a network-based model of rabies spread using GPS telemetry data from dogs in urban N’Djamena, Chad. Other models of rabies-spread in which parameters that describe contact heterogeneity were derived from telemetry data include canine (20) and raccoon models (21, 22).

Patterns of contacts are likely to be altered by the behavioural effects of clinical rabies. Although Hirsch et al. (21) demonstrated that seasonal patterns of rabies incidence in raccoons could be explained by changes in social structure due to normal seasonal behavioural change of the hosts, the influence of rabies-induced behavioural changes on social structure has neither been researched nor explicitly incorporated in simulation models in any species.

Here, our objective was to investigate rabies persistence in small populations of free-roaming dogs – such as those found in rural communities in PNG and northern Australia – by predicting the probability, size and duration of rabies outbreaks. We also investigate the influence of rabies-induced behavioural changes on model predictions and the effect of pre-emptive vaccination on rabies spread in such populations.

## 2. Methods

### 2.1 Disease model and study overview

We developed a stochastic model to simulate social networks of free-roaming domestic dogs and the subsequent transmission of rabies between individual dogs within these networks following the latent infection of a single, randomly-assigned dog. The structure of the social networks was based on three empirically-derived networks of spatio-temporal associations between free-roaming domestic dogs in three Torres Strait Island communities; Kubin, Warraber and Saibai (12). The course of rabies infection in a susceptible dog was simulated in daily time-steps and followed an SEI_1_I_2_R process (rabies infection status: susceptible [S], latent [E], pre-clinical infectious [I_1_], clinical [I_2_] and dead [R]).

Rabies virus transmission from an infectious (*i*) to a susceptible (*s*) dog is described by Equation 1, in which the daily probability of contact between a pair of such dogs was calculated based on the edge-weight between the pair (*E*_*is*_), which is the proportion of a 24 hour period during which that pair of dogs is spatio-temporally associated (in the event of no network connection, *E*_*is*_ = 0). Transmission of rabies further depends on the probability of a bite (*P*_*i*_) by the infected dog conditional on its infection status (I_1_ or I_2_), and the probability of subsequent infection of the susceptible dog (*T*_*s*_). Generation of the social network and estimation of the parameters associated with the dog population dynamics and rabies epidemiology are described in supplementary material S1 and S2, respectively, and parameter values are shown in Table 1. Maximum iteration duration was 3 years.

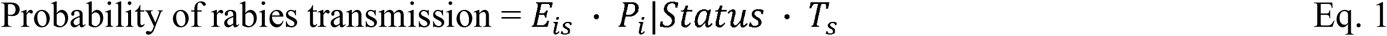

**Table 1.** Parameters used to describe canine rabies epidemiology in a simulation model of rabies spread in a dog population, to determine the influence of social structure on outbreak duration and number of rabies-infected dogs. sd = standard deviation, GSA = global sensitivity analysis. *rabies-induced behavioural changes

| Parameter | Distribution | Parameters | GSA value range | Source |
| --- | --- | --- | --- | --- |
| Population size | uniform | range = 50—90 | 50—90 | Empirical data (pers comm.: |
| Daily per capita birth/death rate | uniform | range = $0.8 \times 10^{-3}$—$1.2 \times 10^{-3}$ | $0.8 \times 10^{-3}$—$1.2 \times 10^{-3}$ | Hudson et al., 2018; field-data, 2016 |
| Incubation period (days) | Lognormal | $\mu = 2.76$-3.13, $\delta = 0.54$-0.75 | 7.4—65.8 days | Tojinbara et al., 2016 |
| Pre-clinical infectious period (days) | gamma | shape = 0.89-3.79, rate = 0.20-1.16 | 0.1—5.7 days | Fekadu et al., 1982 |
| Clinical period (days) | gamma | shape = 2.49-3.56, rate = 0.81-1.19 | 1.1—10.2 days | Hampson et al., 2009 |
| Probability develops furious form | uniform | range = 0.2—0.5 | 0—0.5 | Vaughn et al., 1965; Fekadu and Shaddock, 1984; Fekadu, 1988; Foggin, 1988; Jayakumar et al., 1990; Butler, 1998 |
| Probability of infection following bite | uniform | range = 0.4—0.52 | 0.4—0.52 | Hampson et al., 2009 |
| Bite probability (pre-clinical and dumb) | uniform | range = $0.69 \times 10^{-2}$—$0.69 \times 10^{-1}$ | $0.69 \times 10^{-2}$—$0.69 \times 10^{-1}$ | Empirical data (field study 2016; *unpublished*) |
| *Bite probability (furious) | uniform | range = $0.69 \times 10^{-1}$—0.69 | $0.69 \times 10^{-1}$—0.69 | Authors' assumption, Brookes et al., 2018b, Hampson et al., 2009 |
| Initial 're-wiring' probability, $\beta$ | empirical | Community specific | 0.38—0.44 | See Table 1, re-wiring probability, $\beta$ |
| *Probability of 're-wiring' | uniform | range = 0.05—0.25 | 0—0.25 | Authors' assumption |
| Degree distribution | empirical | Community specific | 5—22 | See Table 1, simulated degree |
| Edge-weight distribution (spatio-temporal association) | empirical | Community specific | $4.1 \times 10^{-7}$—0.42 | See Table 1, simulated edge-weight |
| *Proportional increase in edge-weight during the clinical phase | uniform | range = 0—0.8 | 0—0.8 | Authors' assumption |

Model outputs included distributions of the predicted duration of outbreaks (defined as the number of days from the introduction of a latently infected dog to the day on which infected dogs were no longer present), the total number of rabies-infected dogs during the outbreak and the effective reproductive number, *R*_*e*_, during the first month following incursion (mean number of dogs infected by all dogs that were infected during the first month).

Initially, rabies was simulated in each of the three community networks and the predicted outputs from each model were compared between each other. Statistical tests were used to determine the number of iterations required to achieve convergence of output summary statistics (supplementary material, S3). Global sensitivity analysis using the Sobol’ method (S4) was used to investigate the relative influence of all input parameters on model outputs. To observe the influence of rabies-induced behavioural changes, model outputs from simulations of rabies spread in each of the three community networks with and without parameters associated with rabies-induced behavioural changes were compared. Finally, the impact of pre-emptive vaccination was investigated by randomly assigning rabies immunity to a proportion of the population (10—90%; tested in 10% increments) prior to incursion of the rabies-infected dog in each iteration.

## 3. Results

### 3.1 Community simulations

Each community simulation comprised 10,000 iterations (more than sufficient to achieve convergence of summary output statistics without limiting computational time [Figure S1]).

Predicted outputs are shown in Table 2. The proportion of iterations in which a second dog became infected was greater than 50% in Kubin and Warraber communities, and 43% in Saibai. In these iterations, predicted median and upper 95% duration of outbreaks were longest in Warraber and shortest in Saibai (median: 140 and 78 days; 95% upper range 473 and 360 days, respectively). In the Warraber simulations, 0.001% iterations reached the model duration limit of 1095 days. The number of infected dogs was reflected by the *R*_*e*_ estimates in the first month: 1.73 (95% range 0—6.0), 2.50 (95% range 1.0—7.0) and 3.23 (95% range 1.0—8.0) in Saibai, Kubin and Warraber communities, respectively. The rate of cases during these outbreaks was 2.4 cases/month (95% range 0.6—7.6), 2.0 cases/month (95% range 0.4—6.5) and 2.6 cases/month (95% range 0.5—8.0) in Saibai, Kubin and Warraber communities, respectively.

**Table 2:**
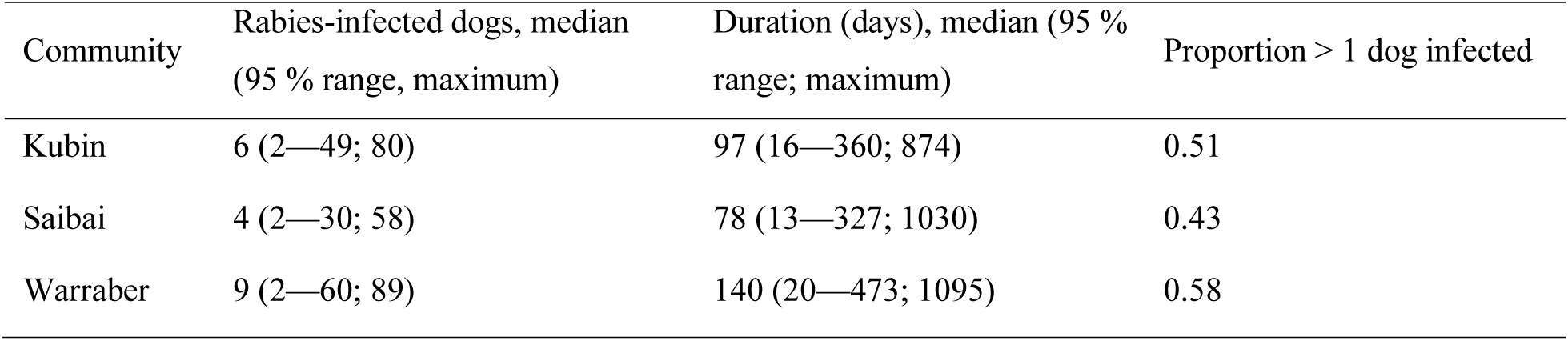
Predicted outputs of rabies spread in incursions in which > 1 dog was infected in simulated networks of dog populations (size 50—90) based on the empirical networks of spatio-temporal association of dogs in three island communities in the Torres Strait, Australia.

### 3.2 Global sensitivity analysis

Figure 1 shows plots of the Sobol’ total-effect sensitivity indices (SI) of parameters for outbreak duration, number of infected dogs and the monthly effective reproductive ratio *R*_*e*_.

**Figure 1.**
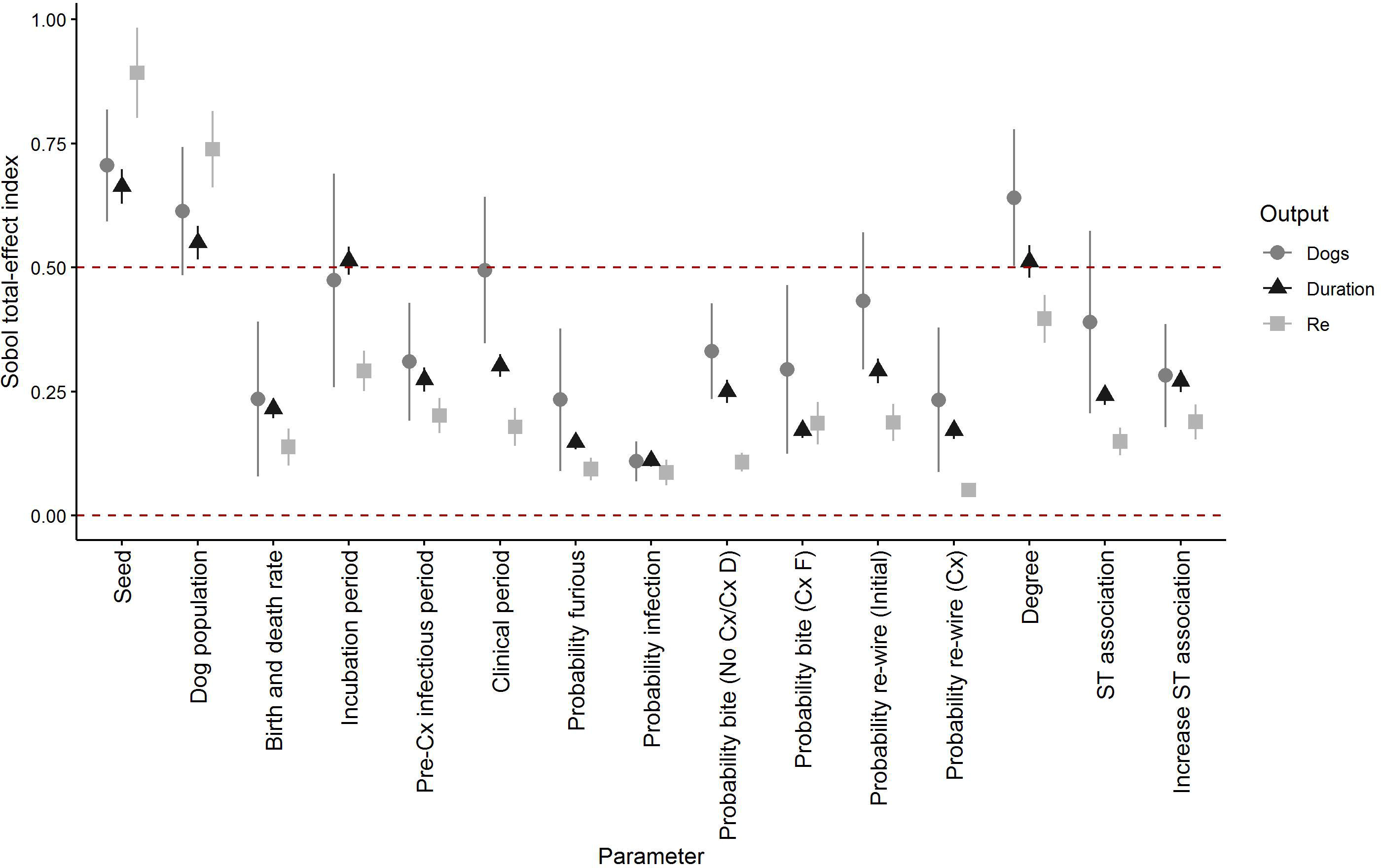
Total effect Sobol’ indices of parameters’ influence on the duration, number of rabies-infected dogs and mean monthly effective reproductive ratio of predicted rabies outbreaks following incursions of rabies in a simulated networks of dog populations based on the empirical networks of spatio-temporal association of dogs in three island communities in the Torres Strait, Australia. Cx = clinical signs, ST = spatio-temporal. Bars indicate 95% confidence intervals.

Supplementary Figure S2 shows Sobol’ first-order effect Sis, which are low relative to the total-effect SIs for all outcomes. This indicates that interactions between parameters are highly influential on output variance in this model and therefore, we focus on the influence of parameters through their total-effects. As expected, the total-effect SI of the seed was highest – it was associated with > 50 % of the variance for all outcomes – because it determines the random value selected in the Bernoulli processes that provide stochasticity to all parameters. The influence of the seed is not presented further in these results.

Incubation period, the size of the dog population and the degree of connectivity were highly influential on outbreak duration (total-effect SI 0.51, 0.55 and 0.51, respectively). All parameters were influential on the predicted number of rabid dogs (total effect SIs > 0.1), and the size of the dog population, incubation and clinical periods, and degree had greatest influence (total effect SIs > 0.5). Dog population size and degree of association were most influential on predicted mean monthly *R*_*e*_ (total effect SI 0.74 and 0.40, respectively).

Of the community-specific parameters (population size, degree and edge-weight distributions, birth and death rates, and initial probability of re-wiring), dog population size and the degree consistently had the greatest influence on each predicted output’s variance. Of network parameters other than degree, the probability of wandering (‘re-wiring’) during the clinical phase (furious form) was markedly less influential than initial ‘re-wiring’ (total effect SIs 0.051 and 0.19, respectively) or either parameter associated with spatio-temporal association (edge-weight; both total effect SIs > 0.15) on predicted mean monthly *R*_*e*_. The influence of the increased probability of a bite by a dog in the clinical period (furious form) on predicted mean monthly *R*_*e*_ was greater compared to the pre-clinical or clinical (dumb-form) bite probability (total-effect SI 0.19 relative to 0.11). The size of the relative influence of these parameters on outbreak duration or number of rabies-infected dogs was reversed and less marked. Birth and death rate consistently had a moderate influence on all outputs (total-effect SI 0.20—0.24).

### 3.3 Rabies-induced behavioural changes

The proportion of outbreaks in which > 1 dog became infected, and the duration, number of infected dogs and *R*_*e*_ in the first month following incursion in simulations without all or with combinations of parameters for rabies-induced behavioural changes, are shown in Figure 2. Outputs from the simulation in each community with all parameters (increased bite probability [furious form], increased spatio-temporal association [edge-weight; dumb form], wandering [‘re-wiring’; furious form]) are included for comparison.

**Figure 2.**
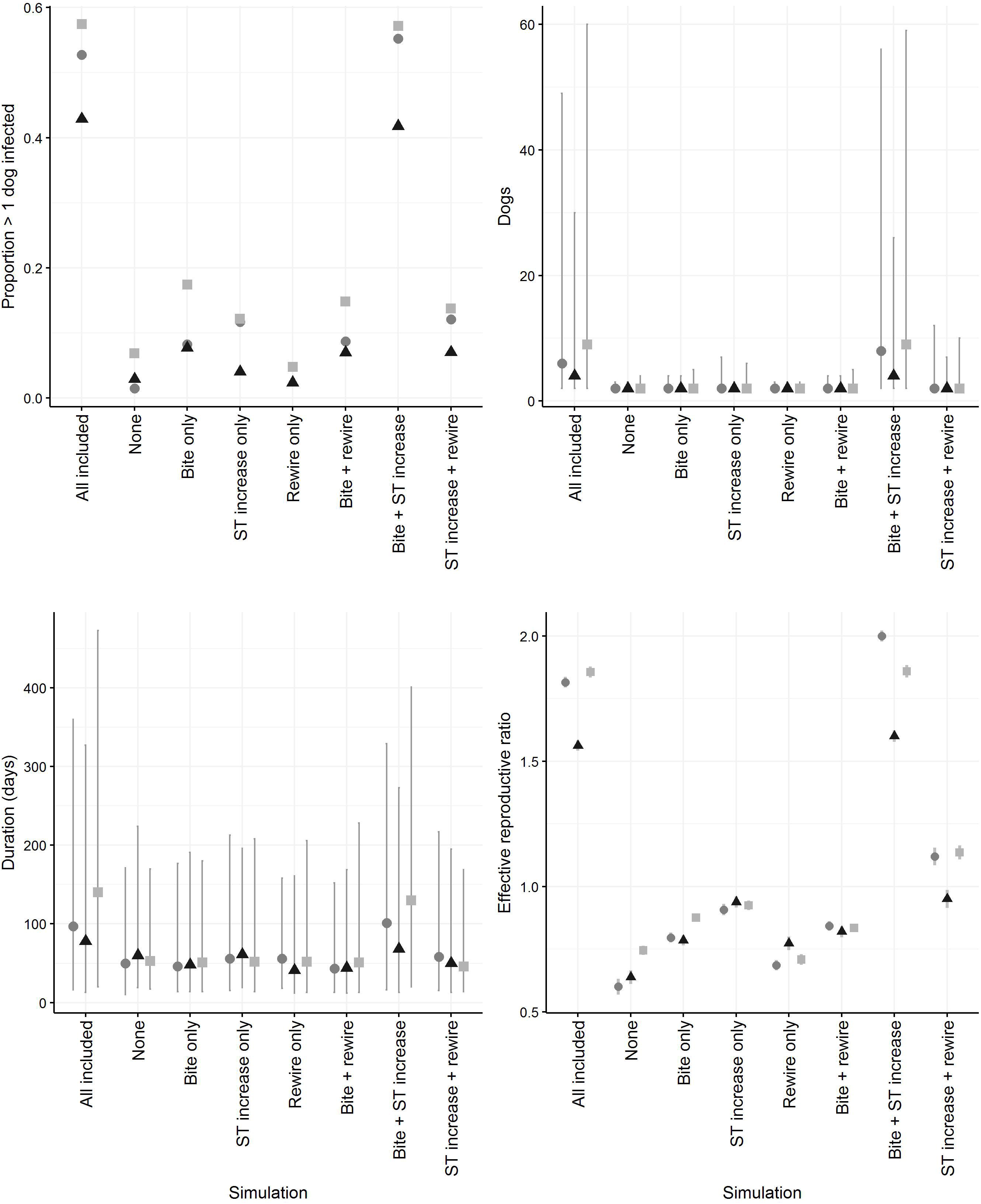
Proportion of predicted outbreaks in which > 1 dog was infected, and the number of infected dogs, outbreak duration and effective reproductive ratio in the first month following incursion, in a simulation model of rabies spread in networks of dog populations based on the empirical social networks of dogs in island communities in the Torres Strait, Australia. Simulations test the influence of inclusion of parameters associated with rabies-induced behavioural change (increased bite probability [associated with the furious form], increased spatio-temporal association [edge-weight; dumb form], wandering [‘re-wiring’; furious form]). Circle = Kubin, triangle = Saibai, square = Warraber. ST = spatio-temporal association. Grey lines = 95 % range.

The simulation without parameters for rabies-induced behavioural changes (Figure 2; ‘None’) propagated following < 10% of incursions in all communities. In 95% of these predicted outbreaks, rabies spread to ≤ 3 other dogs over a median duration of ≤ 60 days. This was reflected by the low *R*_*e*_ in the first month of these incursions (≤ 0.75).

Inclusion of one parameter associated with rabies-induced behavioural changes was still insufficient for sustained predicted outbreaks. Overall, < 20% incursions in these simulations resulted in rabies spread to ≤ 6 other dogs over a median duration of ≤ 56 days. *R*_*e*_ in the first month of these incursions indicated that increased spatio-temporal association, followed by an increased probability of bite were more likely to result in rabies spread than ‘re-wiring’ to increase network contacts in these simulations. This pattern was reflected in the upper 95% range of dogs infected, which was greatest when increased spatio-temporal association was included, and least when ‘re-wiring’ was included.

When combinations of rabies-induced behavioural changes were included, increased bite probability and spatio-temporal association together were sufficient to achieve similar proportions of predicted outbreaks in which > 1 dog was infected (40—60% of incursions) as the simulation with all parameters included (Figure 2 ‘Full’). Predicted impacts and *R*_*e*_ in the first month following incursion were also similar. *R*_*e*_ was greater than the sum of *R*_*e*_ from scenarios with increased bite probability and spatio-temporal association alone.

With combined spatio-temporal association and ‘re-wiring’, the 95% range of the number of infected dogs was greater than simulations in which only one parameter was included (up to 11 other dogs) but *R*_*e*_ in the first month following incursion was close to 1 in all communities, reflecting overall limited rabies spread. In the combined increased bite probability and ‘re-wiring’ simulation, propagation did not occur to > 4 dogs, reflecting the *R*_*e*_ of ≤ 0.8.

### 3.4 The impact of pre-emptive vaccination

Due to the similarity between median outputs from each community and greatest variation in outputs from Warraber, only Warraber was included in this section. All parameters were initially included, then due to observed results, vaccination simulations were also run with a stable population (births and deaths were excluded). Figure 3 shows all outputs.

**Figure 3.**
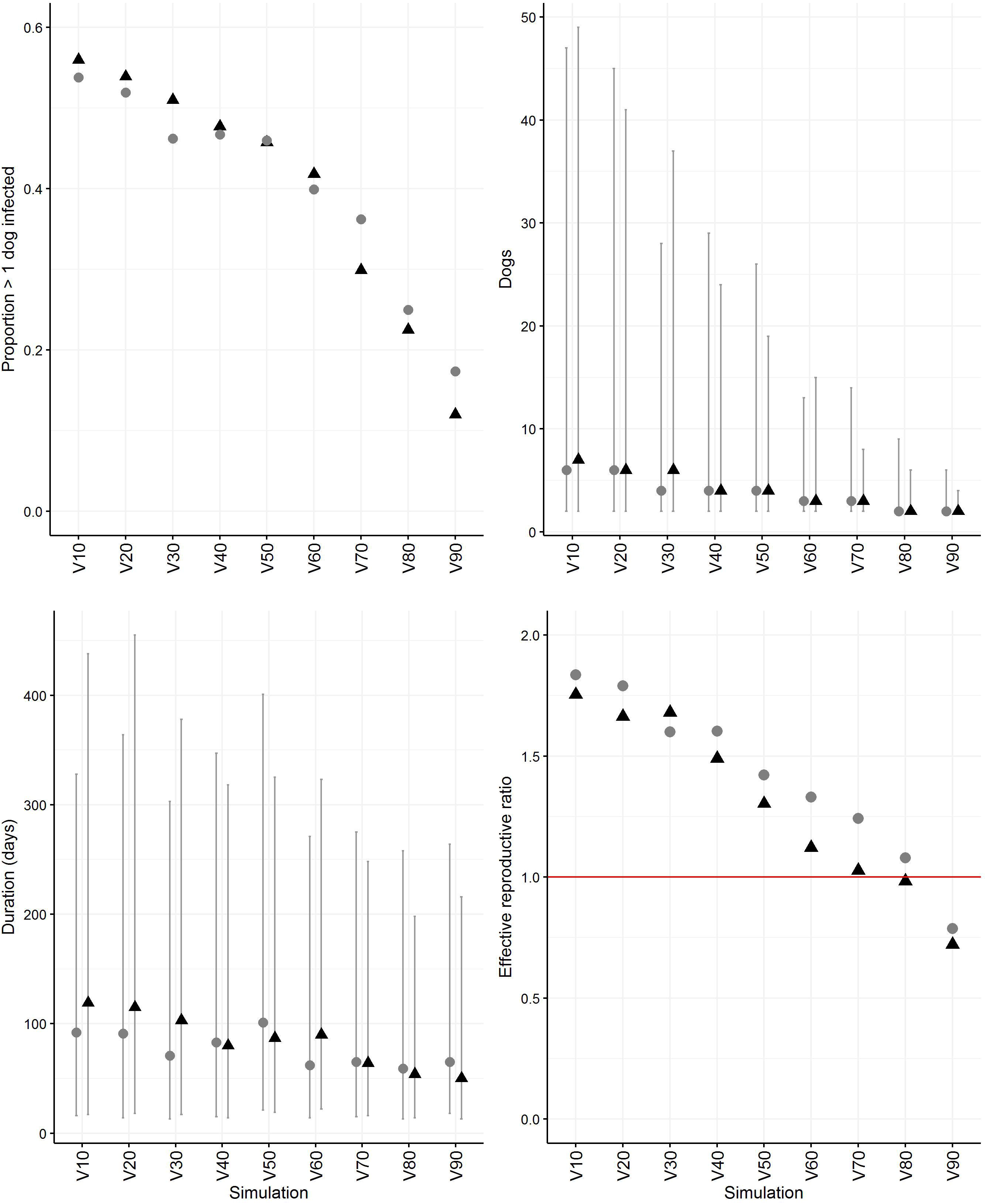
Proportion of predicted outbreaks in which > 1 dog was infected, and the number of infected dogs, outbreak duration and effective reproductive ratio in the first month following incursion in which >1 dog was infected, in a network simulation model of rabies spread based on the empirical social networks of dogs in Warraber, Torres Strait, Australia. Simulations test the influence of pre-emptive vaccination of increasing proportions of the population. Grey circles = simulations with births and deaths included, black triangles = simulations with births and deaths excluded. Grey lines = 95 % range.

In all simulations, the predicted proportion of outbreaks in which > 1 dog was infected fell as the proportion of pre-emptively vaccinated dogs increased – a greater reduction was observed in the simulations with stable populations – and was < 40% when at least 70% of the population were vaccinated. In simulations in which births and deaths were included, the predicted proportion of outbreaks in which more than one dog was infected was still 17% and 12 % when 90% of the population were vaccinated in simulations with and without births and deaths, respectively.

In outbreaks in which > 1 dog was infected, the duration of outbreaks decreased as vaccination proportion increased (although the 95% range was always predicted > 195 days in all simulations). The median number of infected dogs was ≤ 3 once at least 60% of dogs were vaccinated in all simulations, but the 95% range was not consistently < 10 dogs until 80% and 70 % of the population was vaccinated in simulations with and without births and deaths, respectively.

The case rate was median 1.6 cases/month (95% range 0.4—4.6 cases/month) when 70% of the population was vaccinated in simulations with births and deaths, with a median predicted duration of 68 days (95% range 16—276 days). In simulations without births and deaths, the case rate was 1.4 cases/month (95% range 0.4—4.3 cases/month) when 70% of the population, with a median predicted duration of 64 days (95% range 16—248 days). Predicted *R*_*e*_ in the first month following incursion reflected these outputs. At ≥ 70% pre-emptive vaccination, *R*_*e*_ was approximately 1 or less when births and deaths were excluded. However, in the simulations with births and deaths *R*_*e*_ did not fall below 1 until > 80 % of the population were pre-emptively vaccinated.

## 4. Discussion

Our study is unique in that we modelled rabies spread in small populations of free-roaming dogs and incorporated the effect of rabies-induced behavioural changes. Key findings included the long duration of rabies persistence at low incidence in these populations, and the potential for outbreaks even with high levels of pre-emptive vaccination. This has implications for canine rabies surveillance, elimination and incursion prevention strategies, not only in rural areas with small communities, but also for elimination programs in urban areas. We discuss our findings and their implications below.

Without behavioural change, we could not demonstrate rabies propagation in the social networks in the current study; disruption of social contacts appears to be key for rabies maintenance in small populations of dogs. Social network studies have shown that dogs form contact-dense clusters (12, 19). Increased bite probability and spatio-temporal association between contacts (edge-weight in the model) were most influential in our model, but it is possible that ‘re-wiring’ of dogs is also influential in larger populations in which there is a greater probability that a dog would ‘re-wire’ to completely new set of dogs in another cluster, thus increasing total contacts and enhancing spread (degree was also found to be highly influential on rabies spread). Limiting types of behavioural change to particular rabies forms was a simplification for modelling that allowed us to differentiate the effects of types of network disruption. In reality, the association between rabies forms and behavioural changes is likely to be less distinct (24) and thus, rabies spread in small populations could be further enhanced if dogs display a range of clinical signs.

Incubation period also strongly influenced outbreak size and duration, and together with rabies-induced behavioural changes that enabled transmission, is likely to have resulted in the ‘slow-burn’ style of outbreaks (low incidence over long duration) that were predicted by this model. In iterations in which propagation occurred, case rate was generally < 3 cases/month without vaccination, and 1.5 cases/month when 70% of dogs were pre-emptively vaccinated. At such low incidence, we believe that canine rabies is likely to have a low probability of detection in communities where there is high population turnover and aggressive free-roaming dogs can be normal (25, 26). In these populations, dog deaths and fights between dogs are not unusual. Undetected, slow-burn outbreaks in previously free regions are a great risk to humans because rabies awareness is likely to be low. They also provide more opportunity for latently infected dogs to travel between communities either by themselves, or with people, which could result in regional endemicity. Townsend et al (34) suggest a case detection rate of at least 5% (preferably 10%) is required to assess rabies freedom following control measures; surveillance capacity in rabies-free regions such as Oceania should be evaluated and enhanced if required.

Pre-emptive vaccination is another option to protect rabies-free regions; for example, an ‘immune-belt,’ an area in which dogs must be vaccinated, was established in the 1950s in northern Malaysia along the Thai border (27). The World Health Organization recommends repeated mass parenteral vaccination of 70% of dog populations to achieve herd immunity (28). Whilst the origin of this recommendation is unclear, it has been accepted for decades – for example, legislation allowed free-roaming of dogs in designated areas if at least 70% of the dog population was vaccinated in New York State in the 1940s (29) – previous modelling studies of pre-emptive vaccination support this threshold (19, 30-32). We found that vaccination with 70% coverage is expected to provide protection to the majority of dogs and that outbreaks were self-limiting. If inter-community dog movements are unusual, the probability of regional spread is unlikely. However, given predicted upper 95% ranges of 8—14 rabies infected dogs for at least 8 months at 70% coverage, we recommend at least 90% coverage to reduce the effective monthly reproductive ration < 1, limit human exposure, and provide a more certain barrier to regional spread.

In places in which movements are not easily restricted such as urban centres in which dog populations are contiguous, our study indicates that comprehensive vaccination coverage is crucial. The political and operational challenges of rabies elimination are well-documented (33), and lack of elimination or subsequent re-emergence is attributed to insufficient vaccination coverage (< 70% dog population overall, patchy coverage or insufficient duration; (32, 34, 35)) and re-introduction of infected dogs (31, 36). Pockets of unvaccinated dogs within well-vaccinated, urban areas could maintain rabies at a low incidence sufficient to re-introduce rabies as surrounding herd immunity wanes. It is also possible that with comprehensive, homogenous 70% coverage, a low incidence of rabies – such as appears possible at 70% vaccination in our study – is sufficient for endemicity in larger populations but is practically undetectable, giving the appearance of elimination. A higher proportion of vaccinated dogs might be required for elimination, and further modelling studies incorporating behavioural change in larger empirical networks are required to test this hypothesis.

Validation of a canine rabies spread model is challenging, not only because variation between model outputs and observed data can arise from many sources, but because rabies surveillance is passive and case ascertainment is notoriously challenging (35), thus undermining comparison of predicted outputs to observed data. However, although outbreak data from small populations of dogs is scarce, observed patterns of disease spread are consistent with those predicted by the current study (37, 38). Global sensitivity analysis indicated that population size (a parameter of reasonable certainty) and degree of connectivity had the greatest influence on duration, size and initial spread; this makes intuitive sense, and as expected, the largest and longest outbreaks were predicted in the Warraber network which had the highest median degree. Degree distribution might vary between populations of free-roaming dogs; however, a study in N’Djaména, Chad, found that the average degree was 9 and 15 (maximum 20 and 64, respectively) in two populations of size 272 and 237 dogs, respectively (19), which is not dissimilar to the Torres Strait dog populations. Reassuringly, input parameters about which there was more uncertainty – for example, bite probabilities – were less influential on variation in outputs.

By exploring rabies epidemiology in small populations of free-roaming dogs – in which contact heterogeneity was determined in part by their social networks and in part by the disease – our study provides insights into how rabies-induced behavioural changes is important for endemicity of rabies in rural and remote areas. We found that rabies induced behavioural change is crucial for the disease to spread in these populations and enables a low incidence of rabies cases over a long duration. Without movement restrictions, we predict that substantially greater than the recommended 70% vaccination coverage is required to prevent rabies emergence in currently free areas.

## Acknowledgements

This project was funded by the Australian Commonwealth Department of Agriculture and Water Resources. We thank the Torres Strait Island Regional Council, Prescribed Body Corporates, residents and participants in the Torres Strait.

## Supplementary Figures

Figure S1. Convergence test plots using predicted duration of outbreak (days) and number of rabid dogs from a simulation model of rabies spread in free-roaming domestic dogs in Torres Strait communities, Queensland, Australia. a = coefficient of variation (CV) of sets of model simulations of increasing number of iterations. Horizontal red line = 0.025. b = proportion of values of < 0.025 (red line) for the coefficient of variation of the previous 100 sets of simulations for increasing numbers of iterations (x-axis). Horizontal line = 0.95.

Figure S2. First-order effect Sobol’ indices of parameters’ influence on the duration, number of rabies-infected dogs and mean monthly effective reproductive ratio of predicted rabies outbreaks following incursions of rabies in a simulated networks of dog populations based on the empirical networks of spatio-temporal association of dogs in three island communities in the Torres Strait, Australia. Cx = clinical signs, ST = spatio-temporal. Bars indicate 95 % confidence intervals.

## Supplementary Material (one file)

S1. Social network generation description

S2. Parameter estimation

S3. Convergence testing

S4. Sensitivity analysis

## References

1. Taylor LH, Nel LH. Global epidemiology of canine rabies: past, present, and future prospects. Veterinary Medicine: Research and Reports. 2015;6:361–71.

2. Mitmoonpitak C, Tepsumethanon V. Dog rabies in Bangkok. Journal of the Medical Association of Thailand = Chotmaihet thangphaet. 2002;85(1):71–6.

3. Beck S, Gunawardena P, Horton DL, Hicks DJ, Marston DA, Ortiz-Pelaez A, et al. Pathobiological investigation of naturally infected canine rabies cases from Sri Lanka. BMC Vet Res. 2017;13(99):(12 April 2017)-(12 April).

4. Hueffer K, Khatri S, Rideout S, Harris MB, Papke RL, Stokes C, et al. Rabies virus modifies host behaviour through a snake-toxin like region of its glycoprotein that inhibits neurotransmitter receptors in the CNS. Scientific Reports. 2017;7(1)12818.

5. Dobson AP. The Population Biology of Parasite-Induced Changes in Host Behavior. The Quarterly Review of Biology. 1988;63(2):139–65.

6. Amaral AC, Ward MP, Freitas JdC. Estimation of roaming dog populations in Timor Leste. Prev Vet Med. 2014;113(4):608–13.

7. Putra AAG, Hampson K, Girardi J, Hiby E, Knobel D, Mardiana IW, et al. Response to a Rabies Epidemic, Bali, Indonesia, 2008-2011. Emerg Infect Dis. 2013;19(4):648–51.

8. ProMED-mail. Rabies - Malaysia: (Perlis), Canine, OIE, Archive Number: 20150825.3599775, Published Date: 2015-08-25 06:03:38. 2015 Contract No.: 20150825.3599775.

9. ProMED-mail. Rabies - Malaysia: (Sarawak), human, dog bite suspected. Request for information. Archive Number: 20170701.5143911. 2017 06.07.17. Report No.

10. Brookes VJ, Keponge-Yombo A, Thomson D, Ward MP. Risk assessment of the entry of canine-rabies into Papua New Guinea via sea and land routes. Prev Vet Med. 2017;145:49–66.

11. Hudson EG, Brookes VJ, Ward MP. Assessing the Risk of a Canine Rabies Incursion in Northern Australia. Frontiers in veterinary science. 2017;4:141-.

12. Brookes VJ, VanderWaal K, Ward MP. The social networks of free-roaming domestic dogs in island communities in the Torres Strait, Australia. Prev Vet Med. 2018.

13. Craft ME, Volz E, Packer C, Meyers LA. Disease transmission in territorial populations: the small-world network of Serengeti lions. J Royal Soc Interface. 2011;8(59):776–86.

14. VanderWaal K, Gilbertson M, Okanga S, Allan BF, Craft ME. Seasonality and pathogen transmission in pastoral cattle contact networks. Royal Society Open Science. 2017;4(12).

15. VanderWaal KL, Obanda V, Omondi GP, McCowan B, Wang H, Fushing H, et al. The “strength of weak ties” and helminth parasitism in giraffe social networks. Behav Ecol. 2016;27(4):1190–7.

16. Kuperman M, Abramson G. Small world effect in an epidemiological model. Phys Rev Lett. 2001;86(13):2909–12.

17. Keeling M. The implications of network structure for epidemic dynamics. Theor Popul Biol. 2005;67(1):1–8.

18. Craft ME. Infectious disease transmission and contact networks in wildlife and livestock. Philosophical Transactions of the Royal Society B: Biological Sciences. 2015;370(1669):20140107.

19. Laager M, Mbilo C, Madaye EA, Naminou A, Léchenne M, Tschopp A, et al. The importance of dog population contact network structures in rabies transmission. PLoS Negl Trop Dis. 2018;12(8):e0006680.

20. Dürr S, Ward MP. Development of a Novel Rabies Simulation Model for Application in a Non-endemic Environment. PLoS Negl Trop Dis. 2015;9(6):e0003876.

21. Hirsch BT, Reynolds JJH, Gehrt SD, Craft ME. Which mechanisms drive seasonal rabies outbreaks in raccoons? A test using dynamic social network models. J Appl Ecol. 2016;53(3):804–13.

22. Reynolds JJH, Hirsch BT, Gehrt SD, Craft ME. Raccoon contact networks predict seasonal susceptibility to rabies outbreaks and limitations of vaccination. J Anim Ecol. 2015;84(6):1720–31.

23. Brookes VJ, Gill GS, Singh CK, Sandhu BS, Dhand NK, Singh BB, et al. Exploring animal rabies endemicity to inform control programmes in Punjab, India. Zoonoses Public Health. 2018;65(1):E54–E65.

24. Foggin CM. Rabies and rabies-related viruses in Zimbabwe: Historical, virological and ecological aspects. Zimbabwe: University of Zimbabwe; 1988.

25. Brookes VJ, Kennedy E, Dhagapan P, Ward MP. Qualitative Research to Design Sustainable Community-Based Surveillance for Rabies in Northern Australia and Papua New Guinea. Frontiers in Veterinary Science. 2017;4(19).

26. Hudson EG, Brookes VJ, Ward MP. Demographic studies of owned dogs in the Northern Peninsula Area, Australia, to inform population and disease management strategies. Aust Vet J. 2018;96(12):487–94.

27. Tan D, Shukor KbH. The Control of Rabies in Malaysia. Rabies in the Tropics: Springer; 1985. p. 600–3.

28. World Health Organization. WHO expert consultation on rabies: third report. 2018.

29. Korns RF, Zeissig A. Dog, fox, and cattle rabies in New York State; evaluation of vaccination in dogs. Am J Public Health Nations Health. 1948;38(1 Pt 1):50–65.

30. Coleman PG, Dye C. Immunization coverage required to prevent outbreaks of dog rabies. Vaccine. 1996;14(3):185–6.

31. Zinsstag J, Lechenne M, Laager M, Mindekem R, Naïssengar S, Oussiguéré A, et al. Vaccination of dogs in an African city interrupts rabies transmission and reduces human exposure. Sci Transl Med. 2017;9(421):eaaf6984.

32. Townsend SE, Sumantra IP Pudjiatmoko, Bagus GN, Brum E, Cleaveland S, et al. Designing Programs for Eliminating Canine Rabies from Islands: Bali, Indonesia as a Case Study. PLoS Negl Trop Dis. 2013;7(8):e2372.

33. Fahrion AS, Taylor LH, Torres G, Mueller T, Durr S, Knopf L, et al. The Road to Dog Rabies Control and Elimination - What Keeps Us from Moving Faster? Frontiers in Public Health. 2017;5.

34. Cleaveland S, Kaare M, Tiringa P, Mlengeya T, Barrat J. A dog rabies vaccination campaign in rural Africa: impact on the incidence of dog rabies and human dog-bite injuries. Vaccine. 2003;21.

35. Townsend SE, Lembo T, Cleaveland S, Meslin FX, Miranda ME, Putra AAG, et al. Surveillance guidelines for disease elimination: A case study of canine rabies. Comp Immunol Microbiol Infect Dis. 2013;36(3):249–61.

36. Laager M, Mbilo C, Lechenne M, Zinnstag J, Chitnis N. Mathematical modelling of dog rabies transmission in N’Djamena, Chad. Trop Med Int Health. 2017;22:21–2.

37. Butler JRA. The ecology of domestic dogs Canis familiaris in the communal lands of Zimbabwe: University of Zimbabwe; 1998.

38. Tenzin, Sharma B, Dhand NK, Timsina N, Ward MP. Reemergence of Rabies in Chhukha District, Bhutan, 2008. Emerg Infect Dis. 2010;16(12):1925–30.

