## Supplementary figures and images for "Rabies-induced behavioural changes are key to rabies persistence in dog populations: investigation using a network-based model"

### Figure S1

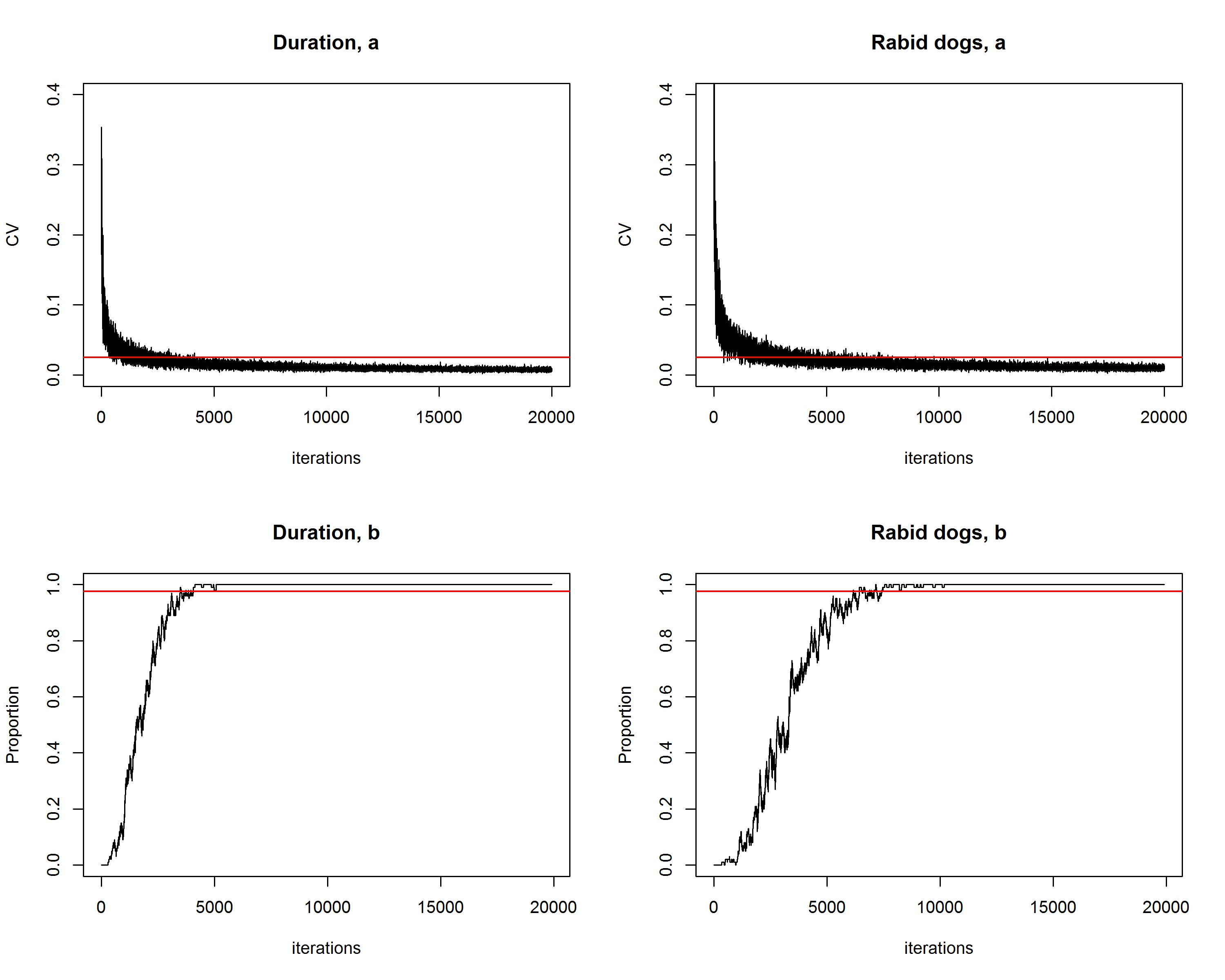

### Figure S2

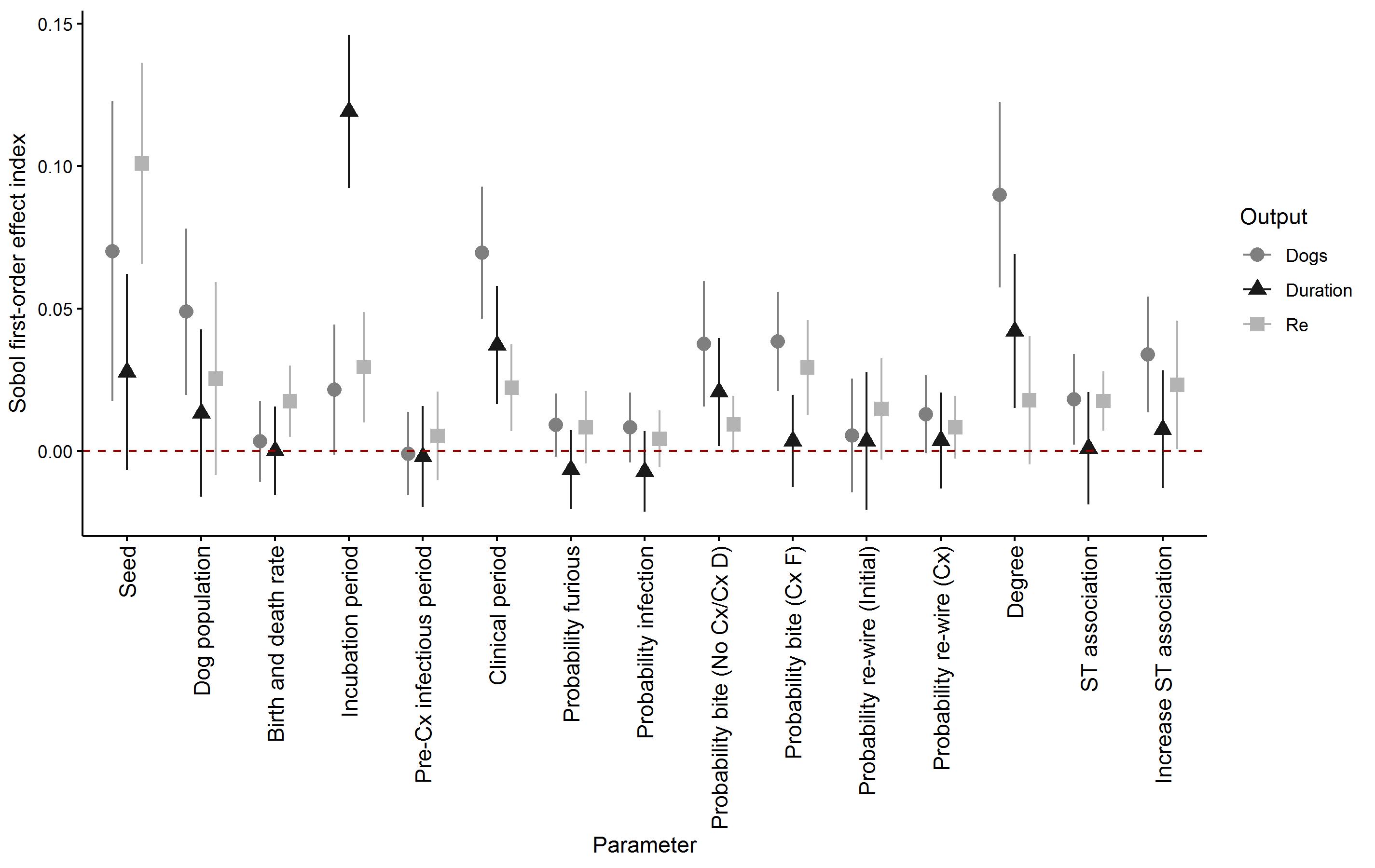
