## Supplementary Material for "Rabies-induced behavioural changes are key to rabies persistence in dog populations: investigation using a network-based model"

**How does rabies spread in small populations of dogs? Investigation using a network-based simulation model**

**Supplementary Material**

***S1	Social network generation***

Prior to each iteration, a modified Watts Strogatz algorithm generated a connected, undirected small-world network of 50—90 dogs with network characteristics that reflected the empirical network, as follows (1-3).

A regular ring lattice was constructed with *N* nodes in which *N* was randomly selected from a uniform distribution of 50—90. Each node was assigned *K* degrees, which was randomly selected from the respective empirical degree distribution of the community represented by the simulation. Each node (*n_i_*) was connected to *K_i_*/2 (rounded to the nearest integer) nearest neighbours in the ring lattice in a forward direction, then all nearest neighbours in a backward direction until *K_i_* was achieved. Existing edges were then re-wired (the edge was disconnected from the nearest neighbour and reconnected to a randomly selected node) following a Bernoulli process (probability *β*) to achieve an average shortest path-length expected in an equivalent-sized Erdõs-Réyni graph in which nodes are connected randomly, whilst maintaining the empirical degree distribution of the community represented by the simulation (4). Edges were then weighted according to the mean expected duration of association between pairs of dogs as a proportion of daily time, and were randomly selected from the respective empirical edge-weight distribution of the community represented by the simulation. Parameters that describe the empirical networks and their derivation are presented in Brookes et al. (5).

Networks simulated with the modified Watts Strogatz algorithm were tested for similarity to the empirical networks prior to use of the algorithm in the model (Table S1). Degree and edge-weight distributions were compared to those of the empirical networks using the Mann-Whitney U and Kolmogorov-Smirnoff tests, to assess similarity of median and shape of simulated distributions, respectively. Mean small-world indices were calculated according to Equation 2 in which *C* is the global clustering coefficient, *L* is the average shortest path length, *s* denotes a simulated network and *r* denotes an Erdos-Reyni random network of equivalent mean degree (6). A small-world index >1 indicates local clustering, consistent with the empirical network structures. Network similarity tests were conducted on 1000 simulated networks for each community.

Small-world Index = ${\frac{C_{s}}{C_{r}}}/{\frac{L_{s}}{L_{r}}}$	Eq. 2

Table S1: Properties of simulated networks of size 50—90 nodes (dogs) and their similarity to empirical networks of spatio-temporal association of dogs, tested by Mann-Whitney U (MWU) and Kolmogorov-Smirnoff (KS), in three island communities in the Torres Strait, Australia., se = standard error.

| Property |  | Kubin | Saibai | Warraber |
| --- | --- | --- | --- | --- |
| Simulated degree | median (range) | 16 (7—22) | 13 (5—16) | 18 (7—20) |
|  | MWU test P value (se) | 0.28 (0.004) | 0.28 (0.004) | 0.29 (0.004) |
|  | KS test P value (se) | 0.82 (0.007) | 0.84 (0.007) | 0.82 (0.007) |
| Simulated edge-weight | median (95 % range) | 0.0004 (0—0.28) | 4.1 x 10^-7^ (0—0.28) | 0.0026 (0—0.42) |
|  | MWU test P value (se) | 0.30 (0.004) | 0.30 (0.004) | 0.32 (0.004) |
|  | KS test P value (se) | 0.83 (0.007) | 0.87 (0.006) | 0.86 (0.006) |
| Re-wiring probability, *β* |  | 0.38 | 0.44 | 0.40 |
| Average shortest path length (se) | | 1.84 (0.002) | 1.80 (0.002) | 1.95 (0.003) |
| Global clustering coefficient (se) | | 0.38 (0.002) | 0.38 (0.001) | 0.31 (0.001) |
| Small world index (se) | | 2.64 (0.01) | 2.61 (0.01) | 2.02 0.01) |

***S2	Parameter estimation***

Parameters that were used to describe the dog populations and rabies epidemiology in the model are listed in Table 1.

***S2.1	Population dynamics***

Population size was selected from a uniform distribution based on the expected size of dog populations in Torres Strait communities (*pers comm*: Torres Strait Island Regional Council and Queensland Health). Births and deaths were included in the model with a daily per capita rate to maintain population size. Given population dynamics of free-roaming domestic dogs in similar communities in the region (7) and field-data from the Torres Strait communities, overall annual population turnover was 30—45% in the model.

***S2.2	Duration of latent, pre-clinical infectious and clinical periods***

A distribution to describe the duration of the pre-clinical infectious period (I_1_) was parameterised by fitting a gamma distribution to data from an experimental study reported by Fekadu et al. (8) in which rabies virus was detected in the saliva of 12 dogs up to 13 days prior to the onset of clinical signs. In the model, the median pre-clinical infectious period was 1.4 days (95 % range 0.1—5.7 days).

We did not find peer-reviewed literature that specifically described the latent period (E). Therefore, a lognormal distribution to describe the incubation period was reproduced from a study in which the date of exposure and onset of clinical signs were extracted from 92 case-records of rabies-infected dogs between 1948 and 1954 in Japan (9). We included uncertainty in the distribution parameters (mean and standard deviation); in the model the median incubation period was 22.2 days (95 % range 7.4—65.8 days). The latent period was then simulated for each infected dog in the model using a randomly selected incubation period and pre-clinical infectious period.

We reproduced a gamma distribution to describe the duration of the clinical period, derived from a study in which the case-histories of suspected rabies-infected animals were reconstructed (10), and included uncertainty in distribution parameters (shape and rate). In the model, median clinical period was 3.29 days (95 % range 1.1—10.2 days) and all rabies-infected dogs died.

***S2.3	Proportion of dogs that develop furious rabies***

In a study in Thailand, 79% of dogs had furious rabies, and in a study in Zimbabwe, Foggin (11) found that 75% of rabid dogs submitted for laboratory examination showed signs consistent with the furious form. However, the proportion of dogs in which the furious form occurs is likely over-estimated in observational studies – dogs exhibiting furious rabies are more likely to roam and bite and are consequently most likely to be recognised and reported as suspect cases. Experimental studies suggest that the proportion of dogs that develop the furious form is less than 50% (12-14). Therefore, we described the probability of developing the furious form of rabies using a uniform distribution with a conservative range (0.2—0.5).

***S2.4	Probability of a bite and subsequent rabies infection***

We made the assumption that the incidence of bites by dogs in the pre-clinical infectious period or clinical period with the dumb form of rabies would reflect the background dog-bite incidence in the communities. Therefore, we assigned a daily probability of bite by an individual dog in these stages of infection according to a uniform distribution derived from a survey of free-roaming dogs in similar communities in the region (0.69 x 10^-2^—0.69 x 10^-1^*; unpublished data;* fieldwork, Northern Peninsula Area, Queensland). To reflect increased aggression and account for uncertainty about this parameter, the daily probability of a bite by a dog with the furious form of rabies in the clinical period was increased by an order of magnitude (0.69 x 10^-1^—0.69). This higher range is also consistent with the distribution of the duration of the clinical period and the findings in previous studies (15, 16).

***S2.5	Network stability***

Altered behaviour during the clinical period is likely to disrupt network contacts. For example, the progressive paralysis and recumbency of a dog with the dumb form of rabies might increase the duration of association between potential contacts, and the wandering of a dog with furious rabies might change network associations (which would increase the total number of contacts during the clinical period). In the model, network disruption was parameterised using a possibility to increase edge-weight to account for an increased duration of association between a dog with the dumb form and its existing contacts, and the possibility to ‘re-wire’ a furious dog’s network connections to other dogs during each 24 hour period of the clinical period of infection. Distributions for these parameters were wide to reflect uncertainty.

***S2.6	Parameters associated with rabies-induced behavioural changes***

Three of the parameters described in the sections above were specifically associated with rabies-induced behavioural changes (Table 1). Each occurred during the clinical period and were either associated with the furious form (an increased bite probability and changed network associations due to altered mentation and aggression) or the dumb form (a possible increase in the duration of association with existing network contacts due to progressive paralysis).

***S4	Convergence testing***

The number of iterations required to achieve sufficient convergence of summary measures was estimated using the following method.

Key output measures – the number of rabid dogs and the duration of outbreaks (days) – were recorded from 9,999 iterations of the model divided equally between all three communities. Ten sets of simulated outputs of an increasing number of iterations (1--5000) were sampled; for example, ten sets of outputs from 1 iteration, 10 sets of outputs from 2 iterations, 10 sets of outputs from 3 iterations, and so on. The mean number of rabies-infected dogs and outbreak duration was calculated for samples in each set. The coefficient of variation (CV; standard deviation/mean) of these sample means was then calculated for each set. With increasing iterations, the variation in sample mean between sets decreases and the CV approaches zero. The number of iterations was considered sufficient to indicate model output stability when 95% of the previous 100 iteration sizes CV was < 0.025.

***S3	Sensitivity analysis***

Variance-based GSA using the Saltelli method was used to determine which parameters most influenced the variance of outputs and was implemented in this study using the SALib module in Python (17). The sequence of events were: parameter sampling to create a matrix of parameter sets for each iteration (parameter ranges are listed in Table 1), simulation using the parameter sets to obtain model output (duration of outbreaks, the total number of rabies-infected dogs and the mean monthly effective reproductive number, *R_e_*), and estimation of sensitivity indices (SIs) to apportion output variance to each parameter. Mean monthly *R_e_* was used as the output of interest in relation to *R* for Sobol analysis, to remove the strong influence of incubation period on *R_e_* in the first month.

To separate the influence of stochasticity from the variation associated with each parameter, the random seed was also included in the Sobol’ analysis (18). The seed value for each iteration was selected from the parameter set (uniform distribution, 1—100).

First-order and total-effect SIs were estimated for each parameter, representing predicted output variance attributable to each parameter without and with considering interactions with other inputs, respectively. SIs were normalised by total output variance and plotted as centipede plots with intervals representing SI variance. Model output variance is most sensitive to inputs with the highest indices.

**References**

1.	Watts DJ, Strogatz SH. Collective dynamics of 'small-world' networks. Nature. 1998;393(6684):440-2.

2.	Brookes VJ. Modified Watts Strogatz Model. 1.0 ed. Github2018.

3.	Hagberg AA, Schult DA, Swart PJ, editors. Exploring network structure, dynamics, and function using NetworkX. Proceedings of the 7th Python in Science Conference (SciPy2008); 2008; Pasadena, CA USA.

4.	Erdös P, Rényi A. On random graphs, I. Publicationes Mathematicae (Debrecen). 1959;6:290-7.

5.	Brookes VJ, VanderWaal K, Ward MP. The social networks of free-roaming domestic dogs in island communities in the Torres Strait, Australia. Prev Vet Med. 2018.

6.	Humphries MD, Gurney K. Network 'Small-World-Ness': A Quantitative Method for Determining Canonical Network Equivalence. PLoS ONE. 2008;3(4).

7.	Hudson EG, Brookes VJ, Ward MP. Demographic studies of owned dogs in the Northern Peninsula Area, Australia, to inform population and disease management strategies. Aust Vet J. 2018;96(12):487-94.

8.	Fekadu M, Shaddock JH, Baer GM. Excretion of rabies virus in the saliva of dogs. J Infect Dis. 1982;145(5):715-9.

9.	Tojinbara K, Sugiura K, Yamada A, Kakitani I, Kwan NCL, Sugiura K. Estimating the probability distribution of the incubation period for rabies using data from the 1948-1954 rabies epidemic in Tokyo. Prev Vet Med. 2016;123:102-5.

10.	Hampson K, Dushoff J, Cleaveland S, Haydon DT, Kaare M, Packer C. Transmission dynamics and prospects for the elimination of canine rabies. PLoS Biol. 2009;7.

11.	Foggin CM. Rabies and rabies-related viruses in Zimbabwe: Historical, virological and ecological aspects. Zimbabwe: University of Zimbabwe; 1988.

12.	Fekadu M, Shaddock JH. Peripheral distribution of virus in dogs inoculated with 2 strains of rabies virus. Am J Vet Res. 1984;45(4):724-9.

13.	Vaughn JB, Gerhardt P, Newell KW. Excretion of street rabies virus in saliva of dogs. Journal of the American Medical Association. 1965;193(5):363-8.

14.	Jayakumar R, Ramadass P, Nedunchelliyan S. Experimental infection with street rabies virus in dogs. Indian Journal of Animal Health. 1990;29(1):7-12.

15.	Brookes VJ, Gill GS, Singh CK, Sandhu BS, Dhand NK, Singh BB, et al. Exploring animal rabies endemicity to inform control programmes in Punjab, India. Zoonoses Public Health. 2018;65(1):E54-E65.

16.	Hampson K, Dushoff J, Cleaveland S, Haydon DT, Kaare M, Packer C, et al. Transmission Dynamics and Prospects for the Elimination of Canine Rabies. PLoS Biol. 2009;7(3):462-71, e1000053.

17.	Herman J, Usher W. SALib: An open-source Python library for Sensitivity Analysis,. Journal of Open Source Software. 2017;2(9).

18.	Johnstone-Robertson SP, Fleming PJS, Ward MP, Davis SA. Predicted Spatial Spread of Canine Rabies in Australia. PLoS Negl Trop Dis. 2017;11(1).
